# Quantitative trait locus mapping reveals an independent genetic basis for joint divergence in leaf function, life-history, and floral traits between scarlet monkeyflower (*Mimulus cardinalis*) populations

**DOI:** 10.1101/2020.08.16.252916

**Authors:** Thomas C. Nelson, Christopher D. Muir, Angela M. Stathos, Daniel D. Vanderpool, Kayli Anderson, Amy L. Angert, Lila Fishman

**Affiliations:** Division of Biological Sciences, University of Montana, Missoula MT USA; Departments of Botany and Zoology and Biodiversity Research Centre, University of British Columbia, Vancouver BC, Canada; School of Life Sciences, University of Hawai’i, Honolulu HI USA

**Keywords:** *Erythranthe*, leaf economics spectrum, life history, mating system, *Mimulus*, physiology, QTL mapping, tradeoff

## Abstract

**PREMISE:** Across taxa, vegetative and floral traits that vary along a fast-slow life-history axis are often correlated with leaf functional traits arrayed along the leaf economics spectrum, suggesting a constrained set of adaptive trait combinations. Such broad-scale convergence may arise from genetic constraints imposed by pleiotropy (or tight linkage) within species, or from natural selection alone. Understanding the genetic basis of trait syndromes and their components is key to distinguishing these alternatives and predicting evolution in novel environments.

**METHODS:** We used a line-cross approach and quantitative trait locus (QTL) mapping to characterize the genetic basis of twenty leaf functional/physiological, life history, and floral traits in hybrids between annualized and perennial populations of scarlet monkeyflower (*Mimulus cardinalis*).

**RESULTS:** We mapped both single and multi-trait QTLs for life history, leaf function and reproductive traits, but found no evidence of genetic co-ordination across categories. A major QTL for three leaf functional traits (thickness, photosynthetic rate, and stomatal resistance) suggests that a simple shift in leaf anatomy may be key to adaptation to seasonally dry habitats.

**CONCLUSIONS:** Our results suggest that the co-ordination of resource-acquisitive leaf physiological traits with a fast life history and more selfing mating system results from environmental selection rather than functional or genetic constraint. Independent assortment of distinct trait modules, as well as a simple genetic basis to leaf physiological traits associated with drought escape, may facilitate adaptation to changing climates.

## INTRODUCTION

Unrelated plant taxa often converge on suites of correlated traits, such as the floral selfing syndrome (Sicard and Lenhard, 2011) or points along the leaf economics spectrum (Wright et al., 2004). Similar spectra sometimes exist within genera and species, with parallel variation in physiological, morphological, and life-history traits across environmental gradients (Mason and Donovan, 2015; Muir et al., 2016; Fajardo and Siefert, 2018; Anderegg et al., 2018; Sartori et al., 2019; Ji et al., 2019). Consistent trait correlations across taxa may arise from fundamental genetic constraints mediated by developmental and physiological tradeoffs. If so, trait syndromes evolve via alleles with pleiotropic effects on multiple traits (Troth et al., 2018; Martínez-Berdeja et al., 2020), establishing strong genetic correlations that may constrain or accelerate adaptation to novel environmental conditions depending on the direction of multivariate selection. Alternatively, convergent integration of traits across gradients may reflect independent local adaptation to shared environmental factors in the absence of any necessary functional connection (reviewed in Donovan et al., 2011; Agrawal, 2020; Guilherme Pereira and Marais, 2020). In the latter case, trait correlations can be readily broken up by gene flow and recombination, allowing relatively unconstrained adaptation along individual trait axes in response to novel selection pressures.

These different genetic bases for trait associations within species imply that distinct processes could underlie current patterns of diversity and also make divergent predictions about evolutionary responses to environmental change. Therefore, characterizing the individual genetic components of trait syndromes, using approaches such as quantitative trait locus (QTL) or genome-wide association mapping, is crucial to understanding the origins and consequences of adaptive covariation among traits.

Along with life history and phenology, leaf functional traits are key mediators of plant adaptive and plastic responses to climatic and geographic variation. The leaf economics spectrum (LES) posits that resource tradeoffs constrain plants to fall along a fast-slow axis of resource gain and expenditure; taxa on one end of the spectrum invest in long-lived leaves with high leaf mass per area (LMA) but relatively low rates of carbon assimilation (photosynthetic rates) and expenditure (growth), whereas those on the other produce leaves with faster rates of both metabolism and turnover (Reich et al., 1997; Wright et al., 2004). The relationships between functional traits and fast-slow life-history axis codified in the LES framework is based on robust patterns of phenotypic variation across large taxonomic and geographic scales (Reich et al., 1997; Wright et al., 2004; Díaz et al., 2016) and is correlated with global variation in plant demography (Adler et al., 2014; Salguero-Gómez, 2017). However, it is not clear what ecological and evolutionary processes (e.g., genetic constraint, natural selection, and/or plasticity) generate these broad patterns (reviewed in Donovan et al., 2011; Agrawal, 2020). Correlations of leaf functional traits along the LES may reflect fundamental physiological/structural constraints (Shipley et al., 2006; Vasseur et al., 2012). For example, community-level assays of phenotypic selection and genetic correlations in desert annuals suggest a lack of the (favored) trait combinations that could break LES correlations (Kimball et al., 2013; Angert et al., 2014). On the other hand, intra-specific variation in leaf functional traits often opposes or blurs LES patterns evident at larger taxonomic scales (Derroire et al., 2018; Anderegg et al., 2018), suggesting that different processes may structure intra-specific vs. inter-specific co-variation among leaf traits. Particularly in herbaceous taxa, cross-species and cross-population variation only partially recapitulate the LES, suggesting that leaf level tradeoffs do not underpin community-level associations (Mason and Donovan, 2015; Muir et al., 2016; Ji et al., 2019; Agrawal, 2020). Furthermore, even when intra-generic associations between climate, life history and leaf ecophysiology are strong, they are ambiguous about mechanism, as population structure and correlated selection can generate association without a necessary functional relationship (Agrawal, 2020). Thus, while such patterns likely reflect past selection, they cannot predict evolutionary responses to novel conditions.

The abundant heritable variation in leaf functional traits among plant populations (Ackerly et al., 2000; Geber and Griffen, 2003; Guilherme Pereira and Marais, 2020) often covaries with life history and mating system variation across climatic gradients. For example, leaf functional traits along a resource-acquisition LES axis correlate with life-history (fast-slow axis) and climate (growing season length) across accessions of the model annual plant *Arabidopsis thaliana* (Sartori et al., 2019). Similarly, photosynthetic physiology correlates with mating system across closely-related *Clarkia* species, perhaps because rapid-cycling annuals require greater reproductive assurance in a temporally constrained growing season that also favors a fast life history and resource-acquisitive leaf traits (Mazer et al., 2010). Such cross-population associations could result from genetic correlations caused by pleiotropy (multiple effects of single genes) or tight linkage (Lowry and Willis, 2010). However, the few studies that have investigated the genetics of leaf functional traits find only weak evidence of LES-predicted genetic coordination among them (Muir and Moyle, 2009; Muir et al., 2014; Taylor et al., 2016; Coneva et al., 2017; Coneva and Chitwood, 2018), and there is even less evidence of genetic coordination of physiology or leaf structure with fast-slow traits at whole-plant level (Ivey et al., 2016). However, the genetic architecture of within- and among-population variation in leaf functional and whole plant history or mating system traits remains poorly understood outside a few model systems (reviewed in Guilherme Pereira and Marais, 2020).

Here, we use a line-cross approach and quantitative trait locus (QTL) mapping to examine the genetic architecture of inter-population divergence in leaf function, life-history, and reproductive (mating system) traits in the riparian plant *Mimulus (Erythranthe) cardinalis*. The hummingbird-pollinated *M. cardinalis* (and its close relative *M. lewisii*, which is bee-pollinated) are well-established models for understanding the evolution of pollination syndromes (Hiesey et al., 1971; Bradshaw et al., 1995; Bradshaw and Schemske, 2003; Yuan et al., 2016) and elevational adaptation (Hiesey et al., 1971; Angert, 2006), as well as species barriers (Ramsey et al., 2003; Fishman et al., 2013; Stathos and Fishman, 2014). Physiological and demographic variation across the wide *M. cardinalis* latitudinal range is also unusually well-characterized, making it model for understanding herbaceous plant responses to rapid climate change (Angert et al., 2011; Sheth and Angert, 2016; Muir and Angert, 2017; Sheth and Angert, 2018). Across its latitudinal range from Baja California to Oregon, *M. cardinalis* occurs in flowing-water riparian habitats, which vary substantially in the timing and magnitude of water availability. Like its congener *Mimulus guttatus* (Lowry and Willis, 2010), *M. cardinalis* cannot survive complete dry down during its vegetative growing season (Sheth and Angert, 2018), so latitudinal variation in water availability strongly shapes population differentiation in life history and phenological traits (e.g. growth rate, flowering time, rhizome production), as well as physiological traits (Muir and Angert, 2017). We focus here on variation represented by parental plants sampled from a montane site in the Sierra Nevada Range (South Fork Tuolumne River) and a desert-scrub site in Southern California (West Fork Mohave River); these sites are separated by only ~1/3 of the latitudinal range of *M. cardinalis*, but differ substantially in precipitation/temperature regimes and represent distinct biogeographic clades in population genomic analyses (Nelson et al. unpublished MS).

In order to quantify the genetic architecture of trait correlations, as well as identify genomic regions contributing to divergence in single or multiple traits, we took a multi-step approach. First, we estimated quantitative genetic parameters (broad-sense heritability and genetic covariances) for 20 traits using an inbred line-cross approach with parental and F_1_ (N = 20 each) replicates and segregating F_2_ hybrids (N = 200). This allows comparison of purely environmental trait associations (within the genetically homogenous genotypes) with the combined environmental and genetic correlations in F_2_s. Second, we generated the first intra-specific linkage map for *Mimulus* section *Erythranthe* using high-density gene-capture markers suitable for scaffolding the draft genome sequence of the species into chromosomes (n = 2152 loci, N = 93 F_2_s). Third, we mapped primary QTLs underlying the 20 traits to determine their individual genetic architecture, then screened for secondary QTLs at a lower statistical threshold and asked whether QTLs for multiple traits co-localized. Although 93 F_2_ individuals do not provide high resolution for characterizing the full number and effect size of QTLs for polygenic traits (Beavis, 1994), this approach provides power to identify major or leading QTLs for individual traits and to rule in or out the possibility that any given trait pair is under the control of major pleiotropic or tightly linked loci.

Together, this work provides insight into genetic architecture of individual adaptive traits as well as the origins of functional trait syndromes and provides a platform for predicting how *M. cardinalis* populations can evolutionarily respond to rapidly changing climatic conditions.

## MATERIALS AND METHODS

### System and plant materials

*Mimulus cardinalis* (Phrymaceae), also known as *Erythranthe cardinalis* (Barker et al., 2012), but see (Lowry et al., 2019), is a hummingbird-pollinated herbaceous plant of low elevation riparian habitats in western North America. Across its latitudinal and elevational range from Baja California to Oregon, *M. cardinalis* varies extensively in life history and functional traits (Muir and Angert, 2017; Sheth and Angert, 2018). In particular, plants from seasonally dry southern populations grow faster and have higher photosynthetic rates than those from wetter northern populations in field common gardens, consistent with an evolutionary shift toward more annual life history strategies in Southern habitats where precipitation is both less available on average and less predictable across years (Muir and Angert, 2017). Demographic studies in the wild confirm adaptive divergence in life history, with southern populations experiencing higher annual mortality and greater reproduction at small size than central (Sierran) populations (Sheth and Angert, 2018). We focus here on representative lines from these respective regions, CE10 (a well-characterized inbred line derived from the Carlon population along the South Fork Tuolumne River in the Sierra Nevada Range near Yosemite National Park; Bradshaw and Schemske, 2003) and WFM (West Fork Mohave River; Fishman et al., 2013; Muir and Angert, 2017).

### Growth conditions, phenotyping, and quantitative genetic analyses

To characterize the genetic basis of coordinated physiological and morphological/life-history differences, we crossed CE10 (as dam) with WFM 1.1 (grown from wild-collected seed) to generate F_1_ hybrids, and then hand-self-pollinated a single F_1_ to generate a segregating F_2_ population. In Fall 2013, F_2_ hybrids (N = 200 total) were grown in a greenhouse common garden at the University of Montana along with replicates of the parental lines (WFM 1 generation inbred) and F_1_ hybrids (N = 20 each). Seeds from each genotypic class were sown on wet sand in separate, parafilm-sealed Petri dishes, with seedlings transplanted into 96-cell growth trays after 10 days (treated as germination time, Day 0) and the entire plugs transferred into 4” pots filled with Sunshine Mix #1 at one month after initial sowing. Plants were maintained in randomized arrays with daily bottom-watering, bi-weekly ½ strength fertilization with Peters Professional 20-20-20 fertilizer, moderate temperatures (80/65 day/night), and supplemental lighting to maintain a long-day photoperiod (16h day/8h night).

To estimate plant size and growth rate, we recorded height (mm) at one time point (Day 45 post-germination) and basal stem diameter (BSD) at two time points 3 weeks apart (Day 33 and Day 55 post-germination), from which we calculated adult relative growth rate (RGR = [ln(BSD2) – ln(BSD1)] / (t_2_ – t_1_)). BSD2 and BSD1 are the basal stem diameters at times t_2_ and t_1_, respectively. At the time of height measurement, we also scored whole plants semi-quantitatively (0-2 scale) for stem anthocyanin production (RedStem). Vegetative anthocyanins may mitigate abiotic stresses (Steyn et al., 2002) and anthocyanin polymorphisms are also associated with life history variation in other monkeyflowers (Lowry et al., 2012). For the first flower to open in each plant (beginning Day 52), we recorded days from germination to flower and measured the length of the gynoecium (distance from pedicel to stigma lobes) and androecium (distance from the pedicel to top of tallest anther). These measures of flower length were used to calculate stigma-anther distance, which generally influences autogamous self-pollination ability (Barrett, 2002).

Midday photosynthetic rate (A_area_, μmol CO_2_ m^−2^ s^−1^), stomatal resistance to water vapor (r_sw_ = 1/g_sw_), and instantaneous water-use efficiency (iWUE = A/g_sw_) were measured on a single leaf (ranging from the 3rd-5th node) over a series of five days (Days 44-49). We analyzed stomatal resistance (r_sw_, m^2^ s mol^−1^) rather than conductance because the former was normally distributed, whereas the latter was right-skewed.

Conditions in the leaf chamber were as follows: leaf temperature = 25°C, CO_2_ = 400 ppm, relative humidity = 0.5-0.75, PAR = 400 μmol quanta m^−2^ s^−1^ (close to ambient for greenhouse in midday), and moderate flow rate (300 umol/s). Leaf functional traits (Day 54), including area (cm^2^), fresh (leaf fresh mass, g) and oven-dried (leaf dry mass, g) weights, were recorded for the paired leaf from the same node as that measured for gas exchange. We calculated LMA (Leaf Dry Mass /Area), leaf dry matter content (LDMC; leaf dry mass /leaf fresh mass) and a proxy for leaf thickness (LMA/LDMC or fresh mass/area, μm) (Vile et al., 2005). Approximately one month after the last plant had initiated flowering (Day 107 from germination), we removed plants from their pots and counted all rhizomes (belowground-initiated vegetative shoots) >3cm long. The total aboveground biomass (not including rhizomes) was oven-dried and weighed (biomass, g).

We compared parental and hybrid means and variances, calculated broadsense heritability (H^2^), and estimated the environmental (r_E_) and genetic (r_G_) correlations among traits with H^2^ > 0.20 following (Fishman et al., 2002). For rhizome number, which was very right-skewed in the F_2_s (many zero values, diverse higher values), we log-transformed the raw counts (+ 1) for these calculations (and later QTL mapping). For the leaf functional traits, individual values may be influenced by ambient conditions and ontogeny. To test for such effects, we conducted analyses of variance (ANOVA) on the full dataset for each trait, including sampling date of physiological measures (date) and leaf pair as factors along with genotypic class. Date was a significant factor (all P < 0.005) for all of the leaf traits, but leaf pair did not matter. Therefore, for the calculations of H_2_, r_E_ and r_G_ for leaf traits, we use the residuals from an ANOVA with date as a fixed factor. However, because genotypic classes were evenly distributed across sampling dates, removing this additional (co-)variance had no qualitative or statistical effect on differences among the parental and F_1_ trait means. Therefore, we present un-standardized means and standard errors for each genotypic class in Table 1. We also ran the QTL analyses (below) for the leaf functional traits with both date-standardized and raw values; however, because these analyses were not qualitatively different in the number or location of QTLs, we present the latter analyses here. All phenotypic analyses were performed in JMP 14 (SAS Institute, 2018).

**Table 1.** Means (± 1 SE) for parental lines and hybrids, plus broad-sense heritability (H^2^), for each trait, organized by trait category. Significant differences among the parental and F_1_ classes (n = 22-25 each; Tukey’s Honest Significant Difference tests) are indicated by superscripts. Calculations were performed on transformed data for rhizomes [log (rhizomes + 1] and date-standardized values for the leaf function traits. For trait where the literature predicts a fast-slow axis of trait co-variation, we have bolded the value of the parent with the more resource-acquisitive leaf function and/or fast life-history.

| TRAIT | CE10 | F<sub>1</sub> | WFM | F<sub>2</sub> | H<sup>2</sup> |
| --- | --- | --- | --- | --- | --- |
| **LIFE HISTORY (LH)** |  |  |  |  |  |
| Basal stem diameter 1 (BSD1) | 1.88 ± 0.08<sup>a</sup> | 2.84 ± 0.05<sup>c</sup> | **2.40 ± 0.07<sup>b</sup>** | 2.62 ± 0.03 | 0.54 |
| Basal stem diameter 2 (BSD2) | 4.14 ± 0.11<sup>a</sup> | 5.16 ± 0.13<sup>b</sup> | **5.71 ± 0.09<sup>c</sup>** | 5.06 ± 0.05 | 0.13 |
| Relative growth rate (RGR) | 0.355 ± 0.026<sup>b</sup> | 0.258 ± 0.014<sup>a</sup> | 0.380 ± 0.010<sup>b</sup> | 0.290 ± 0.006 | 0.20 |
| Height (cm) | 41.6 ± 2.4<sup>a</sup> | 80.4 ± 2.8<sup>b</sup> | **70.2 ± 4.0<sup>b</sup>** | 74.4 ± 1.7 | 0.61 |
| Days to flower | **59.1 ± 0.6<sup>b</sup>** | 54.5 ± 0.5<sup>a</sup> | 63.1 ± 0.8<sup>c</sup> | 56.9 ± 0.3 | 0.41 |
| Rhizomes | 4.76 ± 0.51<sup>c</sup> | 3.08 ± 0.37<sup>b</sup> | **0.36 ± 0.13<sup>a</sup>** | 1.96 ± 0.14 | 0.48 |
| Aboveground biomass (g) | 14.48 ± 0.25<sup>a</sup> | 16.25 ± 0.29<sup>b</sup> | **17.49 ± 0.4<sup>c</sup>** | 16.61 ± 0.13 | 0.29 |
| Stem anthocyanin (RedStem) | 0 ± 0<sup>a</sup> | 0.8 ± 0.13<sup>b</sup> | **2.72 ± 0.09<sup>c</sup>** | 1.25 ± 0.07 | 0.76 |
| **LEAF FUNCTION (LFunct)** |  |  |  |  |  |
| Assimilation (A<sub>area</sub>; μmol CO<sub>2</sub> m<sup>-2</sup> s<sup>-1</sup>) | 10.88 ± 0.3<sup>a</sup> | 11.02 ± 0.28<sup>a</sup> | **12.31 ± 0.23<sup>b</sup>** | 11.47 ± 0.11 | 0.25 |
| Stomatal resistance (r<sub>sw</sub>; m<sup>2</sup> s mol<sup>-1</sup>) | 2.80 ± 0.18<sup>b</sup> | 2.07 ± 0.09<sup>a</sup> | **2.16 ± 0.11<sup>a</sup>** | 2.52 ± 0.05 | 0.43 |
| iWUE (mmol CO<sub>2</sub> mol<sup>-1</sup> H<sub>2</sub>O) | 29.50 ± 1.44<sup>b</sup> | 22.52 ± 0.90<sup>a</sup> | 26.34 ± 1.19<sup>a,b</sup> | 28.27 ± 0.51 | 0.42 |
| Leaf wet mass (g) | 0.58 ± 0.025<sup>a</sup> | 0.64 ± 0.02<sup>a</sup> | **0.79 ± 0.024<sup>b</sup>** | 0.65 ± 0.013 | 0.60 |
| Leaf dry mass (g) | 0.077 ± 0.004<sup>a</sup> | 0.092 ± 0.003<sup>b</sup> | **0.095 ± 0.003<sup>b</sup>** | 0.09 ± 0.002 | 0.58 |
| Leaf area (cm<sup>2</sup>) | 27.36 ± 1.00 | 26.99 ± 0.77 | 28.86 ± 0.89 | 27.39 ± 0.49 | 0.58 |
| Leaf thickness (μm) | **211.1 ± 2.7<sup>a</sup>** | 237.0 ± 2.6<sup>b</sup> | 274.4 ± 4.1<sup>c</sup> | 237.2 ± 1.25 | 0.20 |
| LMA (g m<sup>-2</sup>) | **28.53 ± 1.11<sup>a</sup>** | 34.52 ± 0.93<sup>b</sup> | 33.07 ± 0.88<sup>b</sup> | 32.95 ± 0.33 | 0.07 |
| LDMC (%) | 13.5 ± 0.5<sup>b</sup> | 14.6 ± 0.4<sup>b</sup> | **12.1 ± 0.3<sup>a</sup>** | 13.9 ± 0.2 | 0.10 |
| **FLORAL/MATING SYSTEM (Rep)** |  |  |  |  |  |
| Gynoecium length(mm) | 49.16 ± 0.30<sup>c</sup> | 47.75 ± 0.32<sup>b</sup> | **45.87 ± 0.35<sup>a</sup>** | 46.21 ± 0.13 | 0.29 |
| Androecium length (mm) | 45.52 ± 0.37<sup>b</sup> | 46.58 ± 0.37<sup>b</sup> | **43.87 ± 0.26<sup>a</sup>** | 44.98 ± 0.17 | 0.47 |
| Stigma-anther separation (mm) | 3.64 ± 0.17<sup>c</sup> | 1.17 ± 0.22<sup>a</sup> | **2.00 ± 0.30<sup>b</sup>** | 1.23 ± 0.10 | 0.33 |

### Genotyping and genetic mapping

Plants were genotyped using a custom probe set designed from the *M. lewisii* LF10 v1.1 genome assembly (Yuan, Sagawa, Young, et al., 2013 www.mimubase.org). The probes targeted exons and adjacent sequence of genes 1:1 orthologous to genes in *M. guttatus*, which is segmentally syntenic with *M. lewisii* and *M. cardinalis* despite a different chromosome number (2N = 28 vs. 2N = 16; Fishman et al., 2014). To select genes for enrichment, we extracted all coding sequence from the *M. guttatus* v2.0 annotation (www.Phytozome.jgi.doe.gov) and used BLAST to find unique hits in the *M. lewisii* v1.1 genome (e-value ≤ 10^−10^ and e-value[best hit] – e-value[second-best] ≥ 10^−10^).

Unique hits were queried against the *M. cardinalis* CE10 v1.9 assembly (www.mimubase.org); only sequences present in both *M. lewisii* and *M. cardinalis* assemblies were kept. Lastly, we performed reciprocal best-hit BLAST between the *M. lewisii* and *M. cardinalis* candidate gene sets to filter out close paralogs. The final probe set consisted of 9126 target regions, which all include exon sequence but may also capture neighboring (and potentially more variable) intronic and untranslated regions.

Sequencing libraries were prepared following (Meyer and Kircher, 2010) and using the SeqCap EZ library preparation kit (Roche Nimblegen, Madison, WI, USA). Briefly, genomic DNA was sheared to ~300 bp using a Covaris E220 UltraSonicator (Covaris Inc. Woburn, MA, USA), Illumina adaptors were added via blunt-end ligation, and each library was dual-indexed before multiplexing all libraries into a single reaction for target enrichment. Target enrichment was performed following probe manufacturer’s protocols. To avoid enrichment of repetitive DNA, we used the universal Developer Reagent C_0_T-1 DNA during probe hybridization. Libraries were sequenced on a single lane of Illumina HiSeq 2500 (PE 125). Raw Illumina reads were quality filtered and trimmed for sequencing adaptors using Trimmomatic (Bolger et al., 2014) and aligned to the v1.9 draft *M. cardinalis genome* (http://mimubase.org) using bwa-mem v0.7.15 (Li and Durbin, 2009). Alignments were filtered for minimum quality scores of 29 using samtools v1.3 (Li et al., 2009). We then removed potential PCR duplicates and realigned around indels using Picard Tools (http://broadinstitute.github.io/picard) and called single nucleotide variants with GATK (v3.3-0-g37228af) (McKenna et al., 2010) following GATK best practices.

We constructed a linkage map of exon-capture markers genotyped in a subset of the CE10xWFM F2s (N = 93). The high-density linkage map was generated in Lep-MAP3 (Rastas, 2017), with imputation of missing genotypes. We removed markers with >10% missing genotypes and markers with evidence of strong segregation distortion by χ^2^ test (*p* ≤ 0.001). We assigned markers to linkage groups with a LOD threshold of 13, resulting in eight linkage groups which were ordered by maximum likelihood using the Lep_MAP3 module OrderMarkers2. Lastly, we conservatively removed errors in map order by assigning the same genetic position to markers within 1000 bp of each other (i.e. from the same targeted capture region). Linkage groups were numbered and oriented in accordance with previous *Erythranthe* section maps (Fishman et al., 2013). For QTL mapping, the full dataset was pruned to 720 recombination-informative sites. To identify QTLs, we conducted single interval scans in rQTL2 (scan1, HK method) (Broman et al., 2003), setting trait-specific LOD thresholds for QTL significance with 1000 permutations of the genotype-phenotype matrix. Because we have only moderate power to detect individual QTLs due to low F_2_ sample size, we used a 2-stage approach to identify multi-trait QTLs. First, we identified primary QTLs for each trait significant at experiment-wide α = 0.05 (LOD = 3.54-3.77, depending on trait). We then also identified secondary significant at a less stringent α = 0.15 (overlapping LOD = 3.01-3.12). QTLs were considered co-localized if their 1.5 LOD-drop intervals overlapped. Because QTL coincidence is necessary (but not sufficient) for the inference of pleiotropy, this generous second threshold reduces the chance that we will reject a shared genetic basis for traits due to low power to detect QTLs. For trait combinations whose major QTLs do not co-localize, and which also do not show strong genetic correlation in F_2_s, we can infer that cross-population trait associations reflect the effects of independent genomic regions, plus linkage disequilibrium.

## RESULTS

### Parental differences

In our greenhouse common garden, the parental lines exhibited striking differences in whole plant life-history/growth traits, leaf functional traits, and floral traits associated with mating system (Table 1, Fig 1). The Sierran CE10 line was typically perennial in its growth form, producing abundant rhizomes, whereas Southern WFM1 exhibited annual or “faster” life-history traits. Relative to CE10, WFM was taller and larger in basal stem diameter (BSD) at both time points (Table 1). However, because WFM was larger at the first measurement of BSD (suggesting more early growth or earlier germination), relative growth rates over the 3-week adult measurement interval did not differ.

**Fig. 1.**
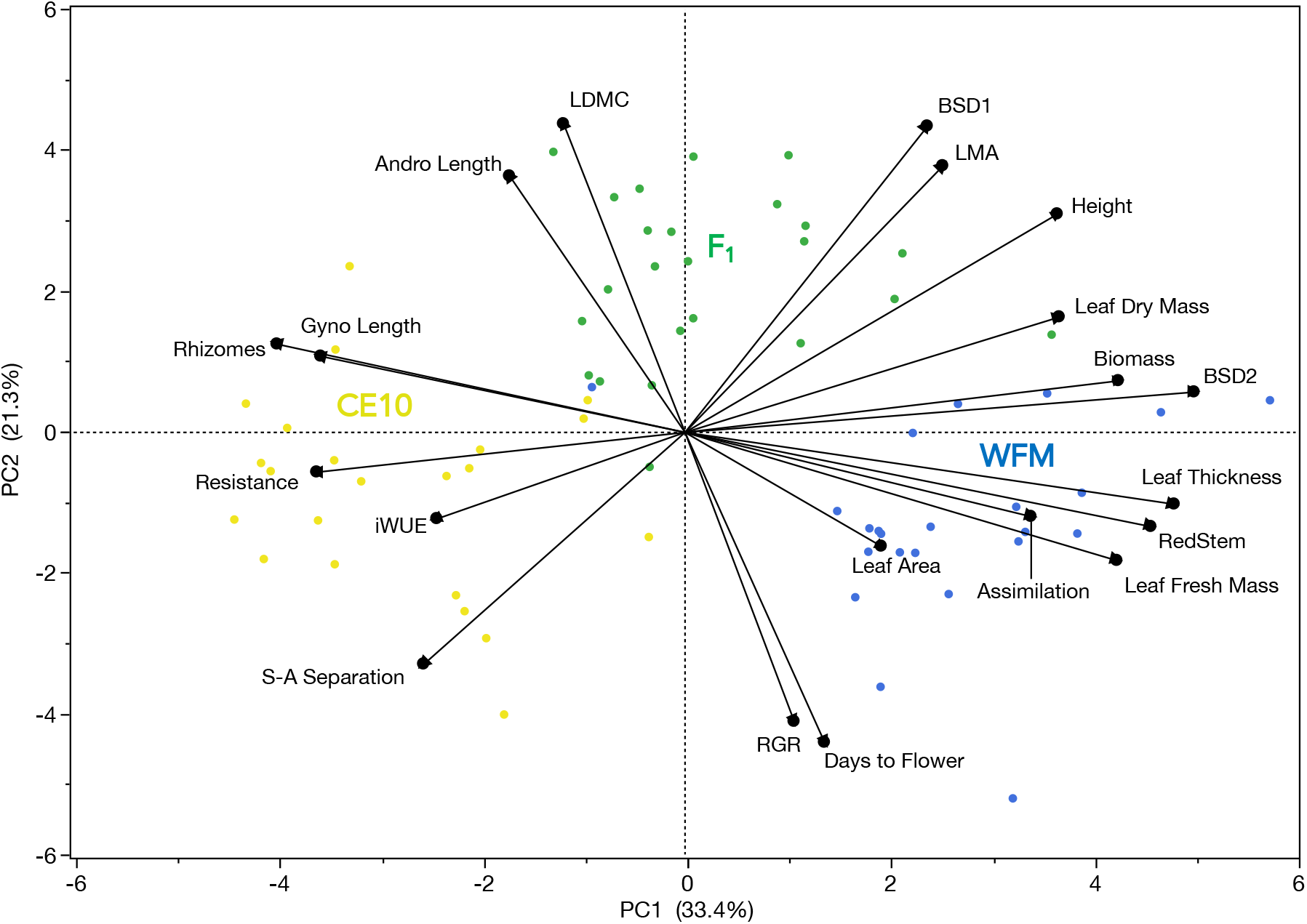
Plot of the first two principal components (PCs) of phenotypic covariation in parental and F_1_ hybrid classes of *Mimulus cardinalis* (points show individuals, colored by genotypic class). PC1 captures the major axis of parental divergence in life history, floral and leaf functional traits, whereas PC2 is loaded by traits exhibiting transgressive variation in F_1_ hybrids.

Most WFM individuals had made no rhizomes by the end of the entire 15-week growth period, instead achieving >20% greater aboveground biomass. The only life-history trait that broke the typical annual-perennial pattern of differentiation was flowering time, with WFM flowering 4 days later than CE10 plants.

Parental lines also differed for physiological and leaf traits consistent with life history adaptation to ephemerally wet (WFM) vs. mesic (CE10) habitats (Table 1). WFM exhibited a strong anthocyanin flush on its stems (mean score 2.72 out of 3), whereas CE10 plants did not flush (all 0s). WFM leaves had higher photosynthetic assimilation rate (A_area_; P < 0.001, Table 1) and lower stomatal resistance to water loss (r_sw_: P = 0.001), consistent with a live-fast, die-young strategy. This translated into lower instantaneous water use efficiency (iWUE) in WFM under well-watered greenhouse conditions. Although the parental lines did not differ significantly in leaf area (P = 0.24), WFM leaves had 36% greater wet mass (P < 0.0001), 25% greater dry mass (P < 0.005), and 15% greater leaf mass per area (LMA; p < 0.005) than CE10 leaves. The greater fresh leaf bulk of WFM vs. CE10 reflects increased water content and/or laminar thickness (P < 0.005) rather than higher tissue density, as WFM actually had 10% lower LDMC (P = 0.01).

WFM had overall smaller flowers than CE10 (> 3.2 mm shorter gynoecium, ~1.6 mm shorter androecium) and significantly lower stigma-anther separation (Table 1), consistent shift towards increased autogamous selfing capacity in plants from the drier and more variable Southern habitat.

### Heritabilities and genetic covariances in hybrids

Most traits differentiated between the parents exhibited overall additivity (F_1_ and F_2_ hybrids, on average, intermediate between parents) or dominance (F_1_ hybrids equivalent to one parent, distinct from the other) (Table 1). However, F_1_s exhibited heterosis (means significantly higher or lower than either parent) for days to flower, basal stem diameter at time 1 (BSD1) and relative growth rate (RGR). F_1_s grew faster at early stages and flower earlier than either parent, then slowed down as adults. In addition, photosynthetic rate and stomatal resistance exhibited opposite patterns of dominance, with F_1_s matching the low parent of each trait and thus having transgressively low water use efficiency (iWUE) (Table 1). Stigma-anther separation was similarly transgressive, with F_1_s having unexpectedly low stigma-anther separation (Table 1).

Broadsense heritabilities (H^2^) were high for the life history traits of flowering date, early growth (BSD1), height, and (log) rhizome number (all >0.40), intermediate for RGR (0.20) and final biomass (0.29), and low for BSD2 (0.13) (Table 1). The essentially Mendelian trait of stem anthocyanin had the highest H^2^ (0.76), but leaf size traits (dry mass, wet mass, and area) were also highly heritable (H^2^ > 0.58). The high heritability for leaf area (despite no parental mean difference) reflects high segregating variance in the F_2_ hybrids for all leaf size traits (including transgressive values well outside the range of the other classes). Leaf physiological traits (photosynthetic rate, stomatal resistance, and iWUE) and leaf thickness, as well as floral traits, all had moderate to high values of H^2^ (0.20-0.50; Table 1). In contrast, the composite LES traits of LMA and LDMC had low H^2^ (< 0.10).

As expected, functionally linked traits (e.g., photosynthetic rate and stomatal resistance) tended to show high genetic (r_G_) and environmental (r_E_) correlations (Fig 2). However, the decomposition of r_G_ and r_E_ also revealed high r_G_ among traits with no environmental correlation in the parental and F_1_ plants (e.g. photosynthetic rate and flowering date). The latter is consistent with genetic correlation due to linkage or pleiotropy, in the absence of any environmental covariance. However, because estimation of r_G_ from F_2_ line cross data requires the assumption of no shift in the covariance matrix or gene x environment or gene x gene (epistatic) interactions, high values for r_G_ do not necessarily imply strong genetic constraints. Even weak linkage among underlying loci in F2s can generate high r_G_ if Cov_E_ (plastic covariation within the genetically invariant classes) is near zero or opposite in direction to the genetic associations. For example, r_G_ is as highly negative for the (functionally unrelated) trait pair of stomatal resistance and stigma-anther separation as it is for resistance and its functional partner photosynthetic rate, likely due to linkage between separate leaf physiology and floral QTLs (see below).

**Fig. 2.**
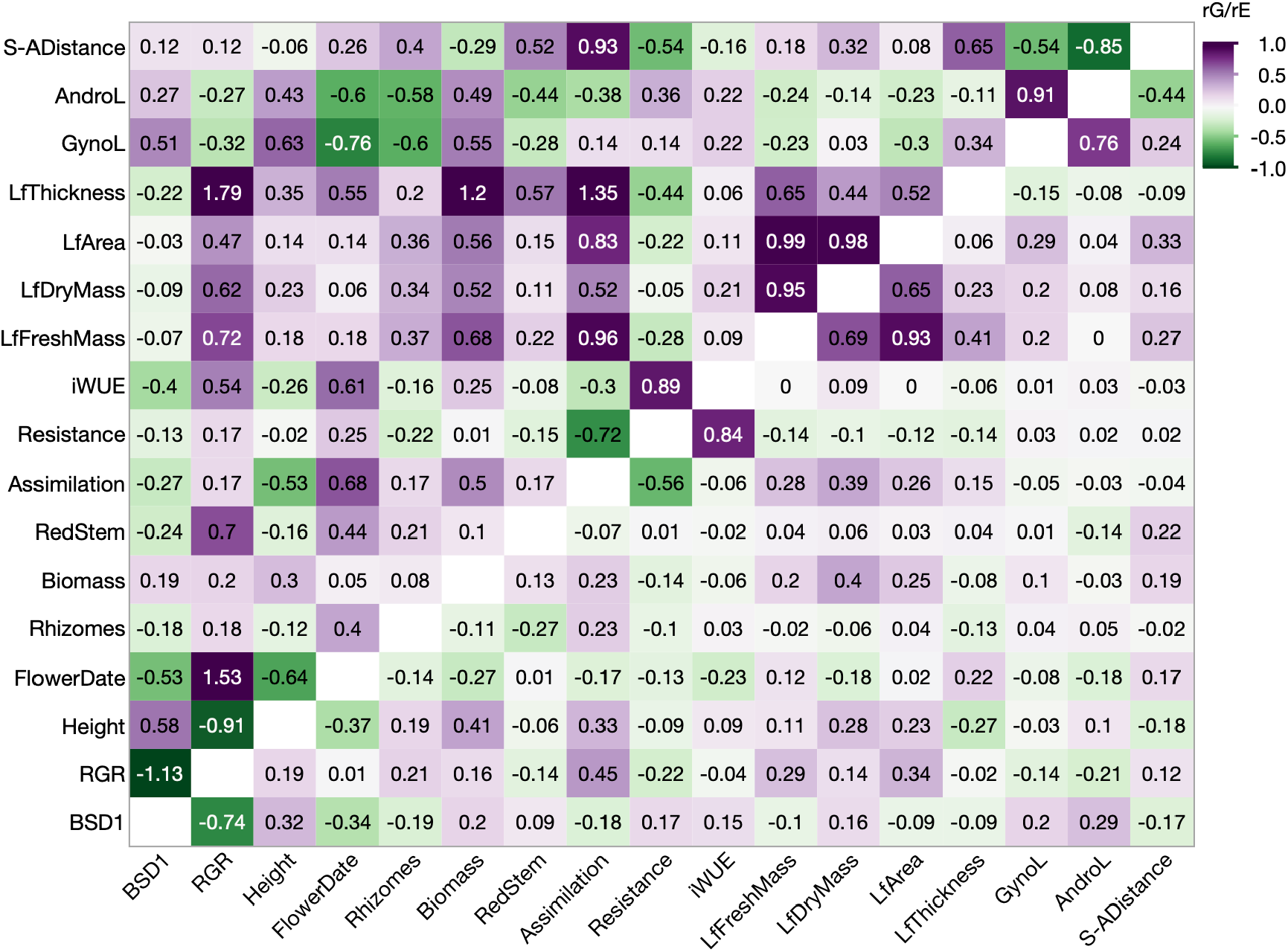
Environmental (r_E_; below the diagonal) and genetic (r_G_; above the diagonal) correlations calculated from parental and hybrid covariances for traits with H^2^ ≥ 0.20. Because r_G_ is calculated assuming r_E_ in the non-recombining classes equals r_E_ the segregating F_2_ hybrids, values can be greater than 1 or < −1, especially when r_E_ is low or opposite in direction to the F_2_ phenotypic covariance.

### Linkage and QTL mapping

The sequence-based genotyping yielded 8100 informative single nucleotide polymorphisms (SNPs) across 2152 gene-capture targets. Genetic mapping resolved the expected 8 linkage groups, with 720 unique (at least 1 recombinant in 186 meioses) marker positions spanning a total of 537 cM (Fig. 3). Overall, the mapped markers exhibited mild but significant (at α = 0.005 level, to control for multiple non-independent tests) transmission ratio distortion in six regions across five of the chromosomes. All of the distorted regions were characterized by excess heterozygosity (>60%), perhaps due to weak selection against homozygotes (inbreeding depression), and two also had a significant deficit of WFM alleles (<42%). In these most extreme cases, on Chromosomes 2 and 4, the frequency of the rarer WFM homozygote was ~10% (9/93 F_2_s) rather than the expected 25%.

**Fig. 3.**
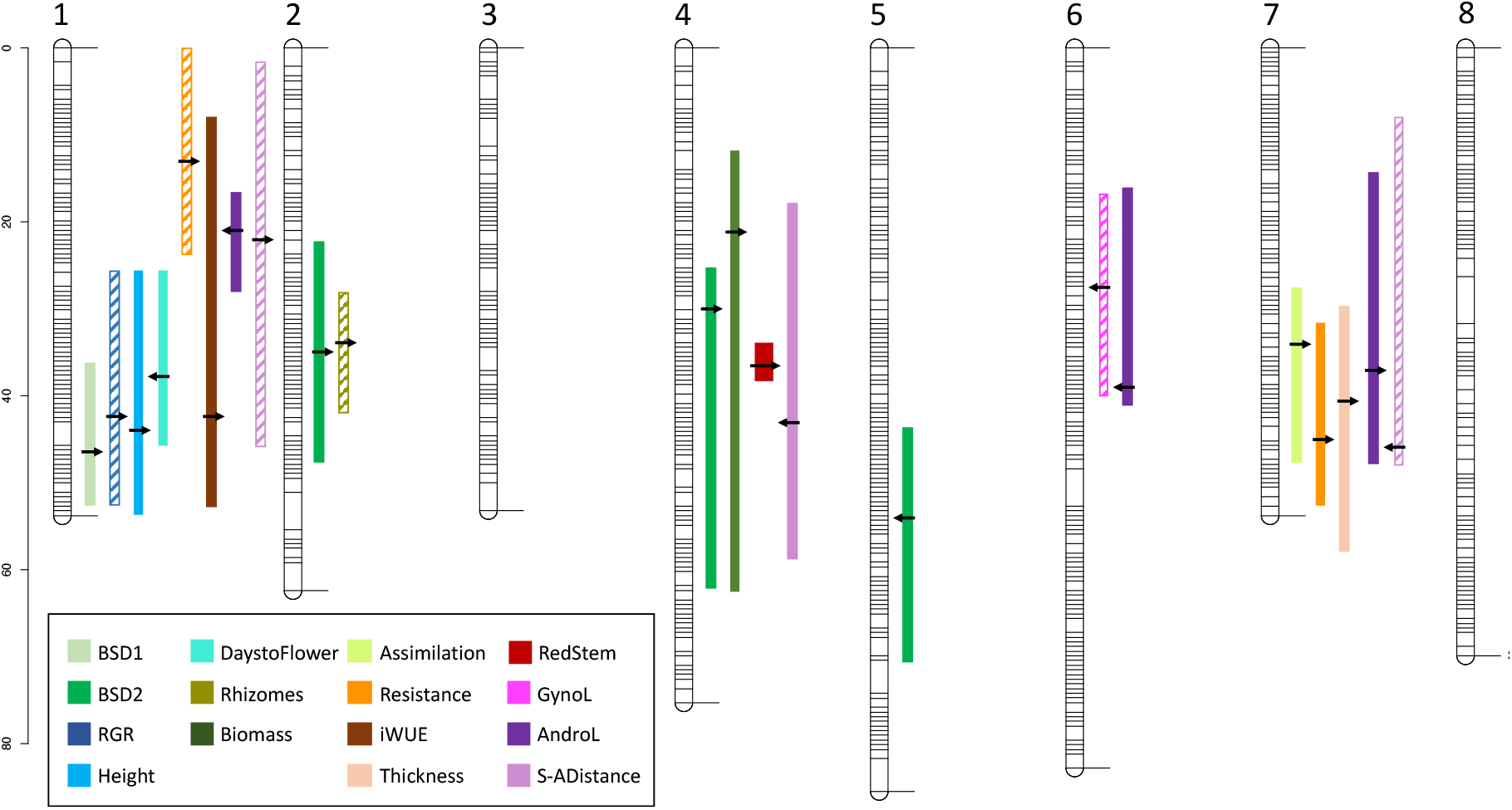
Quantitative trait loci for life history, leaf function, and floral traits in intra-population *Mimulus cardinalis* hybrids, mapped onto the eight linkage groups/chromosomes. Scale at left is centiMorgans (cM), and the positions of 720 linkage-informative gene-based markers are shown on linkage groups. Colored bars (wide = r^2^ > 0.25) show the 1.5 LOD drop confidence interval around each peak (solid: primary QTLs at genome-wide α = 0.05 LOD thresholds 3.40-3.78, depending on trait; hashed: secondary QTLs at genome-wide α = 0.15, LOD > 3.02 - 3.14, depending on trait). Arrows show the location of QTL peaks, with the arrowhead pointing right if the direction of allelic effects matches the expectation from parental means. See Table 2 for QTL groupings and effect sizes.

We detected 16 primary QTLs for 10 traits (genome-wide α = 0.05), and six more overlapping QTLs (at secondary α = 0.15 threshold) for a total of 13 traits (Fig. 3, Table 2). For life-history traits, we detected QTLs in four regions, with all but one affecting multiple traits. On Chromosome 1, a multi-trait life-history QTL (LH1) was significantly associated with BSD1, height, and days to flower, and (less strongly) relative growth rate (but not BSD2). This QTL explains 15-20% of the variance for traits with moderate (0.2 - 0.6) heritability, and Sierran CE10 alleles appear recessive. The allelic effects at LH1 are in the expected direction (e.g., CE10 genotypes growing slowly early and reaching shorter heights) for all traits except flowering time. The three other life history QTLs all underlie variation in BSD2; notably, LH5 (the largest at r^2^ = 0.21) solely affects BSD2 in opposition to the parental difference, while WFM alleles at the two multi-trait QTLs (LH2 and LH5) increase BSD2 while decreasing rhizome number and increasing biomass, respectively (Table 2).

**Table 2.**
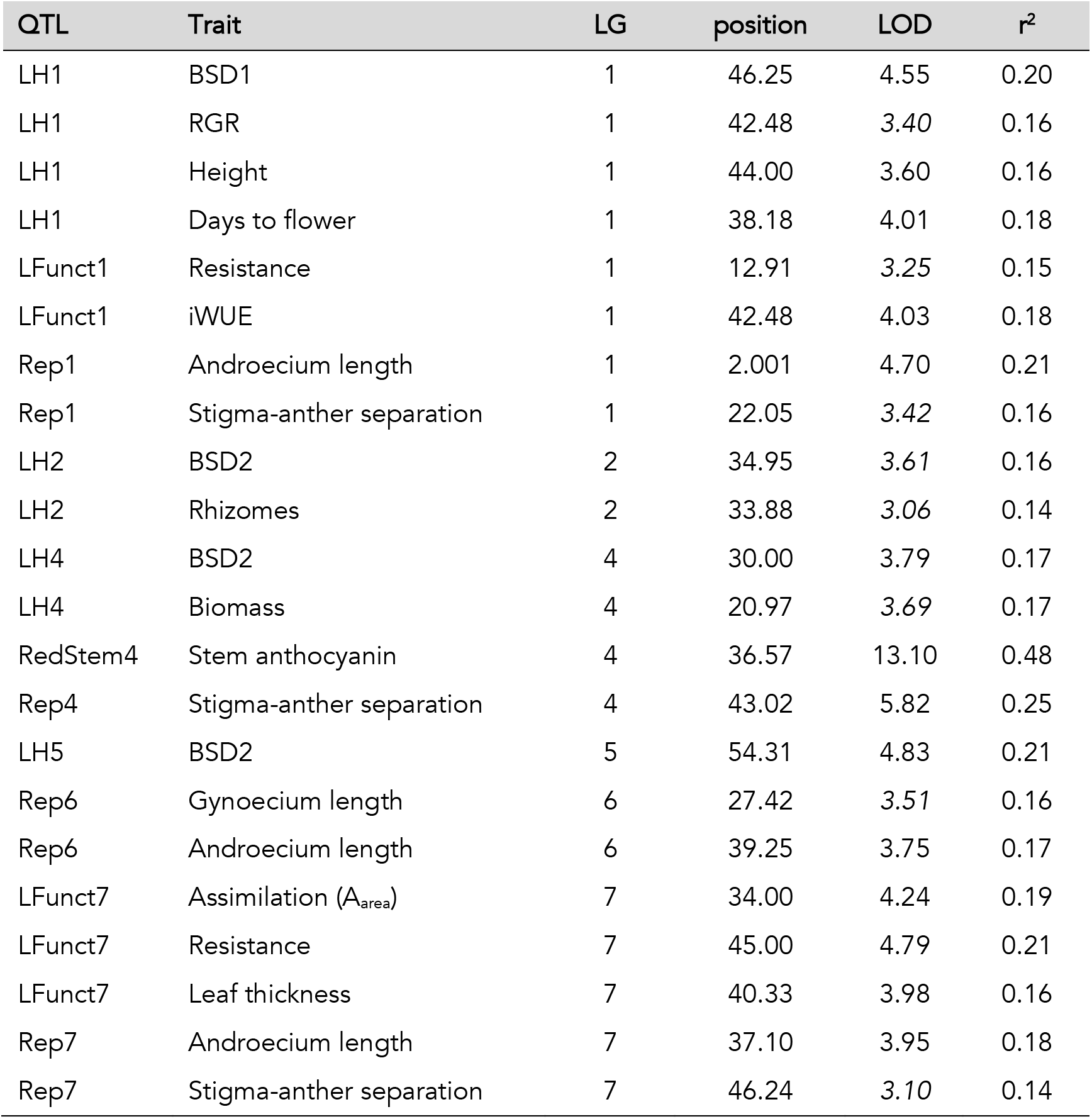
Quantitative trait loci (QTLs), grouped by trait category and chromosome if overlapping within a category, with peak cM position, LOD score, and r2 (percent of F_2_ variance explained) for each trait. LOD scores only significant at the lower secondary threshold (α = 0.15) level are italicized.

For physiological/functional traits, we detected a Mendelian QTL on Chromosome 4 for anthocyanin production on the main stem (RedStem4; max LOD = 13.1, p < 0.00001). RedStem4 had additive effects, with heterozygotes (like F_1_s) intermediate between CE10 (no anthocyanin flush) and WFM (strong flush). This suggests a simple regulatory switch controlling vegetative anthocyanin production under benign greenhouse growth conditions. We detected no QTLs for the three leaf size traits, despite substantial H^2^ (Table 1), or for the low-heritability traits LMA or LDMC. A broad region on Chrom 1 (LFunct1) was associated with F_2_ variation in iWUE, with CE10 alleles recessively increasing water use efficiency.

LFunct1 overlaps with a secondary QTL for stomatal resistance (CE10 homozygotes high, as expected from parental difference), but the region had no significant association with photosynthetic rate. Finally, we mapped a tight cluster of QTLs on Chrom 7 (LFunct7) for photosynthetic assimilation and stomatal resistance (r^2^ = 0.19 and 0.21, respectively), as well as leaf thickness (r^2^ = 0.16). Because the two gas exchange traits have only H^2^ of 0.25-0.43 (i.e., a large fraction of the date-standardized F_2_ phenotypic variance is not genetic), this major QTL may be sufficient to explain much of their between-population genetic differentiation. Indeed, the two homozygous F_2_ genotypes at LFunct7 are more differentiated than the parental means (both standardized and unstandardized values).

Floral traits associated with mating system mapped to QTL regions on four chromosomes. Rep1 affected both androecium length (r^2^ = 0.21, WFM bigger) and stigma-anther separation (r^2^ = 0.16, WFM smaller), while Rep7 had similar effects in the opposite direction (androecium length: r^2^ =0.18, WFM smaller; stigma-anther separation: r^2^ = 0.14, WFM bigger). Rep6 affected gynoecium and androecium length in concert (r^2 =^ 0.16 and 0.17, respectively; CE10 homozygote shorter for both), so had no effect on stigma-anther separation. Rep4 was strongly associated with stigma-anther separation (r^2^ = 0.25), but not (significantly) with the length of male or female reproductive organs separately. The allelic effects of this major QTL go in the opposite direction from the expectation from the parental difference. That is, WFM homozygotes (N = 16, mean = 2.44) have much greater stigma-anther separation than CE10 homozygotes (N = 26, mean = 0.53), with heterozygotes intermediate. The variable directionality and dominance of individual floral QTL effects in hybrids likely contributes to the unexpectedly low stigma-anther distances of F_1_s (Table 1). All three stigma-anther separation QTLs remained significant in a model with all included, but we did not have sufficient F_2_ numbers to scan for epistatic interactions.

## DISCUSSION

### Multi-trait divergence between populations reflects life-history adaptation to local soil moisture regimes

The transition between perenniality and annuality is one of the most common in flowering plants, changes the context for selection on numerous other traits, and yet remains poorly understood from both ecological and genetic perspectives (Lundgren and Marais, 2020). Although previous work has demonstrated latitudinal clines in population demographic variables and common garden phenotypes associated with a fast-slow life history axis (Muir and Angert, 2017; Sheth and Angert, 2018), *M. cardinalis* is generally considered to be a perennial species. Nonetheless, our comparison of Sierran (CE10) and Southern (WFM) parental accessions indicates that the latter are morphologically specialized for an annual lifestyle, with fast early growth and no rhizome production under greenhouse conditions. This divergence parallels the life-history polymorphism in the yellow monkeyflower (*M. guttatus),* in which widespread annual and perennial ecotypes sort by local soil moisture conditions and are defined by chromosomal inversions generating strong associations among co-favored traits (Lowry and Willis, 2010). *M. cardinalis* experiences strong viability selection in its southern range (Sheth and Angert, 2018), particularly severe drought periods when some sub-sites dry down completely in late summer (A. Angert, pers. obs.). In that demographic study, WFM had the lowest survival rate of >30 populations tracked, suggesting ongoing selection for traits that maximize reproduction prior to dry-down. For a strictly annual population, survival to flowering, rather than annual survival, matters for population dynamics; however, Southern *M. cardinalis* populations are not strictly annual and low rates of year-to-year survival were not ecologically compensated for by seedling recruitment (Sheth and Angert, 2018). Nonetheless, WFM plants may be relatively well-adapted to a of highly variable summer water. More work will be necessary to fully characterize phenotypic variation, selection, and evolutionary responses in wild *M. cardinalis*, but Southern populations may have potential to respond evolutionarily to climate shifts (see also Sheth and Angert, 2016) and to share adaptive variation with Northern populations experiencing increased drought (Paul et al., 2011).

The annualized life-history of WFM Southern *M. cardinalis* is accompanied by genetic differences in leaf physiological traits consistent with an adaptive drought-escape strategy (Table 1). Several trends parallel expectations for a fast-slow axis of LES variation: annual, early-big WFM plants appear more “resource-acquisitive” in that they have relatively high area-standardized photosynthetic rates, plus lower stomatal resistance and water use efficiency (Table 1). However, these physiological differences are positively associated with leaf thickness differences between populations. Both LMA and leaf thickness are greater in WFM plants, but LMDC is lower (Table 1), suggesting that WFM’s higher fresh mass reflects disproportionately increased water content per unit area (i.e. succulence or thickness) rather than increased cell density or investment in cell wall. Although not consistent with LES predictions (Wright et al., 2004; Poorter et al., 2009), this combination of high photosynthetic assimilation rate with relatively high LMA in the more arid and seasonal climate parallels patterns in similar herbaceous taxa (Mason and Donovan, 2015). However, even within herbs, leaf functional traits may experience different selection from similar environmental conditions (e.g. for tolerance vs. escape) depending on life history. For example, in perennial *Silene latifolia*, populations from xeric sites tended to be have thicker leaves but also slower growth trajectories compared to mesic populations (Delph, 2019). Similarly, work with annual *Arabidopsis* accessions suggests that complex combinations of environments (perhaps associated with winter vs. spring annual life histories) may be as important as any single environmental axis in shaping joint variation in fast-slow and LES traits (Sartori et al., 2019). Although the finding that within-species LES correlations often run counter to between-species predictions has been attributed to intraspecific plasticity in individual traits (Anderegg et al., 2018), this work and other common garden studies (Muir et al., 2016; Ahrens et al., 2020) suggest that genetic (and likely adaptive) differences within species may also contribute to unexpected outcomes.

In flowering plants, shifts in life history are often associated with shifts in reproductive traits, as the evolution of annuality in ephemeral habitats can precipitate (or at least be correlated with) selection on traits associated with successful pollination (e.g., phenology and mating system) (reviewed in Barrett et al., 1996; Gaudinier and Blackman, 2020). Paralleling latitudinal clines in previous studies of *M. cardinalis* (Muir and Angert, 2017), our southern WFM accession flowers slightly later than the Sierran CE10 line.

Slowness to flower in the faster-growing Southern annual may reflect adaptation to a longer growing season compared to Sierran populations bounded by Spring and Fall freezes. Particularly if hummingbirds appear in Summer at both sites, intrinsic time to flower may be uncoupled from whole plant growth rate. Intriguingly, Southern *M. cardinalis* populations are also uniquely responsive to artificial selection on flowering time (Sheth and Angert, 2016), suggesting that spatially and temporally variable soil moisture may maintain abundant genetic variation for this trait.

Greater environmental unpredictability in Southern populations could also account for the smaller flower size and reduced stigma-anther separation (classic selfing syndrome traits; Sicard and Lenhard, 2011) of WFM *M. cardinalis*. In ephemerally moist soils throughout Southern California, *M. cardinalis* co-occurs (and hybridizes) with the obligately annual tiny-flowered selfer *M. parishii* (Fishman et al., 2015).

Quantitative shifts in stigma-anther separation in Southern *M. cardinalis* likely reflect shared environmental conditions that create pollinator limitation and select for self-fertilization as reproductive assurance (as in *M. guttatus*: Fishman and Willis, 2008; Bodbyl Roels and Kelly, 2011). However, the community of hummingbird pollinators in each region, abiotic factors selecting on correlated corolla traits, or drift/inbreeding could have also led to the difference. Reduced stigma-anther separation (relative to Sierran plants) has also been reported for *M. cardinalis* collected near the Northern range limit (Hazle and Hilliker, 2015), suggesting selection for increased selfing at both range edges. However, more work will necessary to characterize intraspecific floral and mating system variation and its interactions with other phenotypes across the species range.

### Composite functional traits often exhibit transgressive means and low heritability in hybrids, suggesting independent genetic bases for the underlying components

Two patterns seen in our experimental hybrids data underline complexities in predicting the evolution of ecological-proxy composite leaf functional traits. First, due to independent assortment and dominance, F_1_ hybrids between isolated populations may be more extreme that either parental genotype (heterosis or underdominance, respectively) for the composite trait (Rieseberg et al., 2011; Thompson et al., 2019). In our cross, F_1_ hybrids have the low photosynthetic rates of the CE10 parent coupled with the low stomatal resistance of the WFM parent, resulting in transgressively low hybrid iWUE; stigma-anther separation is similarly underdominant (Table1). Such transgressive composite phenotypes may reduce the fitness of hybrids between divergent populations and slow adaptive introgression and evolutionary rescue under contemporary climate change (Thompson et al., 2019). Second, later-generation hybrids may exhibit low heritability or transgressive segregation because genes underlying component trait sort into novel combinations (Rieseberg et al., 1999), potentially with epistatic interactions (Muir and Moyle, 2009). In this study, two composite LES traits (LMA and LDMC) exhibited notably low heritability (< 0.10), despite substantial parental differentiation and high heritability of the underlying component measures (Table 1).

These patterns raise the question of whether composite traits such as LMA and iWUE evolutionarily diverge as direct targets of selection or are generally a by-product of responses to selection on component traits. The answer undoubtedly depends on the particular functional trait and its genetic architecture. For example, the composite floral trait of stigma-anther separation rapidly responds to selection for increased autogamous self-fertilization in *M. guttatus* (Bodbyl Roels and Kelly, 2011) and water use efficiency may similarly be a direct target of selection in *M. cardinalis*. However, other composite traits such as LMA may lack both heritability (as here) and a functional relationship to fitness variation within species. Thus, separating the individual components of composite functional traits may be key to predicting their evolutionary trajectories, especially in novel genetic backgrounds and environments.

### *The first intraspecific genetic map for* Mimulus *section* Erythranthe *provides a key resource for ecological and comparative genomics in this model system*

Genetic mapping in *Mimulus* section *Erythranthe* has focused on inter-specific crosses (Bradshaw et al., 1995; Fishman et al., 2013; 2015) and/or narrow genetic introgressions containing induced mutations or candidate loci for species differences (Yuan, Sagawa, Young, et al., 2013; Yuan, Sagawa, Di Stilio, et al., 2013). However, *Erythranthe* species pairs are distinguished by chromosomal inversions that suppress recombination and translocations that cause inter-chromosomal pairing and severe sterility in hybrids (Fishman et al., 2013; Stathos and Fishman, 2014). Thus, inter-specific maps involving *M. cardinalis* have resolved only 6 or 7 linkage groups (Fishman et al., 2013; 2014; 2015), confounding the genetics of adaptive differentiation with processes of chromosomal divergence and speciation. Our gene-anchored intra-specific map cleanly resolves the expected 8 *Erythranthe* chromosomes to advance study of the comparative evolutionary genetics of trait divergence across *Mimulus*. For example, the Mendelian RedStem4 QTL occurs in the center of *M. cardinalis* Chromosome 4, which is syntenic with *M. guttatus* LG8 (Fishman et al., 2014). A cluster of three R2R3 MYB genes on LG8 has underlies floral and vegetative anthocyanin polymorphisms in the *M. guttatus* species complex (Cooley et al., 2011; Lowry et al., 2012), and the syntenic region also controls corolla anthocyanin patterning traits in section *Erythranthe* (Yuan et al., 2014). This evidence of a conserved regulatory basis for vegetative anthocyanins encourages comparative analyses of stress tolerance and pigmentation. Further, within-*M. cardinalis* genetic mapping of diverse traits provides a framework for future ecological genomics using approaches complicated by linkage disequilibrium (e.g. genome-wide association mapping) or blind to traits (e.g., F_ST_ outlier analysis).

### Multi-trait QTLs suggest modular adaptation, but only weak co-ordination between fast-slow life history and leaf functional traits

QTL analyses reveal little genetic co-ordination between life history and leaf functional traits, despite identification of multi-trait QTLs within each category. The multi-trait LH1 QTL had parallel effects on early growth (BSD1), flowering time, and height (r^2^ =0.13-0.20, CE10 homozygotes slower-growing, shorter, and later to flower), but no impact on final biomass. The coordinated plant architecture effects of LH1 resemble the major DIV1 QTL underlying the perennial to annual transition in yellow monkeyflowers (Hall et al., 2006; Lowry and Willis, 2010; Friedman et al., 2015); however, it had no detectable effects on rhizome production, a key trait for perennials. Overall, fast vs. slow early growth and flowering also appeared decoupled from late growth and final biomass; three additional QTLs (LH2, LH4, and LH6) influenced BSD2, with little or no effect on BSD1. Counter to the mean parental difference, WFM alleles in LH2 and LH4 regions increased BSD2, while also decreasing rhizome production (LH2) and increasing final biomass (LH4), respectively. This pattern suggests strong modularity of early and late growth, with the former genetically associated with flowering phenology and the latter mediating total aboveground biomass and potentially trading off against rhizome production. The molecular genetic basis of perennial to annual transitions remain surprisingly opaque across herbaceous plants (reviewed in Friedman and Rubin, 2015), and this evidence of modularity suggests that *M. cardinalis* may be a tractable system for exploring the mechanistic basis of within-species variation in life history. LH1 is our best (but still modest) candidate for co-incidence of life history and leaf functional QTLs, as overlaps with a QTL affecting iWUE through elevated stomatal resistance in CC plants. LH1/LFunct1 effects are consistent with fast-slow axis predictions, with “slow” CE10 alleles relatively resource-conservative in terms of leaf traits (high iWUE and low stomatal resistance). However, although the iWUE and RGR peaks are co-localized the iWUE QTL is extremely broad (Fig. 3); therefore, tight genetic linkage or pleiotropy are unlikely explanations for LH1/LFunct1 co-localization.

Importantly, the largest leaf functional QTL (LFunct7) has no significant effects on growth traits under common garden conditions (Table 2). Instead, this genomic region coordinately influences multiple aspects of leaf structure/physiology, with WFM alleles additively increasing photosynthetic rate (A_area_) and leaf thickness while reducing stomatal resistance (r_sw_). It is not surprising that photosynthetic rate and resistance are strongly (negatively) correlated; one obvious source of pleiotropy would be differences in the density or regulation of stomata, with WFM exchanging more CO_2_ (and losing more H_2_O) over the same area. However, both populations are amphistomatous (stomata on both upper and lower leaf surfaces), and do not differ in their stomatal density (202 mm^−2^ for CE10 versus 200 mm^2^ for WFM, C. Muir, unpublished data). Amphistomy is expected for taxa from high-light habitats, such as *Mimulus*, and may allow for greater delivery of CO_2_ to mesophyll cells for a given leaf thickness, enhancing the efficiency of photosynthesis when light is not limiting (Mott et al., 2006; Muir, 2019). Although laminar thickness is difficult to measure directly on large numbers of plants (Vile et al., 2005), anatomical work suggests that thicker leaves generally have taller mesophyll palisade cells and greater photosynthetic efficiency (Terashima et al., 2001). Therefore, the co-localization of gas exchange and leaf thickness QTLs in this study is interesting from both a functional and an evolutionary perspective, as all three traits may be pleiotropic effects of variation in the thickness of the mesophyll layer. Plants with thick or succulent leaves, like those with high LMA, are generally considered to be on the slow/resource-conservative end of the LES (Wright et al., 2004). Consistent with that idea, two previous genetic studies (in tomato and *Arabidopsi*s) have found that leaf thickness is genetically associated with slower growth, and this has been interpreted as evidence of a leaf-level tradeoff (Coneva et al., 2017; Coneva and Chitwood, 2018). In contrast, our WFM plants combine alleles for rapid early growth with unlinked variation supporting that growth with high photosynthetic rates in relatively thick leaves. This suggests that the broad patterns seen across taxa and climates (Adler et al., 2014; Salguero-Gómez, 2017) may not derive from genetic or resource constraints at the leaf-level, at least not over the scale of population divergence (Agrawal, 2020).

More broadly, our results suggest that the ratio of leaf wet mass to area (thickness) may be a better predictor for area-standardized photosynthetic capacity (and a fast life history) than LMA, at least in high-light-adapted taxa. Succulence is often associated with drought tolerance in arid habitats, but here greater leaf thickness is genetically associated with fast, resource-acquisitive, drought-escape strategy in a light-loving herb. A slightly thicker, more succulent leaf could have more chloroplasts and mesophyll surface area per leaf area, increasing light-saturated photosynthetic rate (Nobel, 2009). In *Poplar,* trees from shorter growing seasons have leaves with higher photosynthesis rates genetically associated with anatomical differences increasing the exposure of mesophyll cell surface area to airspace (Soolanayakanahally et al., 2009; Milla-Moreno et al., 2016). LFunct7 suggests this may be a common physiological mechanism under simple genetic control between *M. cardinalis* populations. However, further work on variation in leaf functional traits within species, particularly those spanning perennial and annual strategies, may reveal whether this is a general axis of plant adaptation across environmental gradients.

The major effects of the LFunct7 QTL, as well as LH1 (affecting flowering, early growth, and plant architecture), make them appealing candidates for further mechanistic exploration. Higher resolution approaches will be necessary to determine whether or not these multi-trait QTLs represent single loci with broad pleiotropic effects, adaptively suppressed recombination among genes for traits under correlated selection (as in the annual-perennial supergene DIV1 in M. guttatus; Lowry and Willis, 2010), or linkage that will be broken up by additional generations of recombination. However, even if they resolve into numerous genes, these genomic regions may be key contributors to the extensive photosynthetic and life history variation across *M. cardinalis’* latitudinal range (Muir and Angert, 2017), facilitating life-history and physiological adaptation to seasonally dry habitats within its Southern range. Furthermore, the substantial (and potentially pleiotropic) effects of these QTLs suggest that genome-wide analyses of natural variation will successfully identify major adaptive loci segregating across the landscape, and that rapid contemporary adaptation may be possible (with gene flow) as climates throughout the species range become more variable.

### Caveats and limitations

Although we had power to detect major QTLs and rule out joint (major) effects of a given genomic region on multiple traits, our small F_2_ mapping population prevents strong inferences about the absolute size and number of loci affecting quantitative traits. Any QTLs above a statistical threshold may show an upward bias in effect-sizes (Beavis, 1994), and we empirically did not have power to detect QTLs explaining less than 14% of the F_2_ variance (Table 2). Indeed, for several divergent and heritable traits (e.g., rhizomes, leaf fresh and dry mass, gynoecium length), no QTLs were significant at our genome-wide statistical threshold (permutation-based α = 0.05). This suggests the action of numerous small-effect loci on these traits (or substantial epistasis); minor loci undoubtedly also influence those traits for which we did map QTLs. In addition, our two-step approach to identifying coincident QTLs could generate bias toward finding an excess of apparent multi-trait QTLs; however, only two additional QTLs would have been identified (one for stigma-anther separation, one for leaf thickness) if we had used the lower secondary threshold (α = 0.15) in our initial genome-wide scan. A small mapping population also reduces power to finely map QTLs and distinguish tight linkage or pleiotropy from simple chromosome-sharing. However, it is notable that none of the leaf functional traits were highly genetically correlated with BSD1 (early growth), flowering time, or rhizome production, suggesting that our QTL-based inference of no tight coordination between leaf physiology and these aspects of the fast-slow axis is robust. Further, numerous multi-trait QTLs suggest that we also had power to detect any strong genetic associations between functional and life history QTLs Finally, our mapping focused on a single pair of populations, represented by an inbred line from the Sierras and the (inbred) progeny of a single accession from Southern California. This means that we are not capturing the full *M. cardinalis* phenotype-genotype map and that inbreeding depression may contribute to patterns of phenotypic variation, particularly for traits (such as stigma-anther separation) where we detected QTLs with effects opposite to the parental mean difference. However, our parental differences in physiology parallel those seen in previous surveys of the species entire range (Muir and Angert, 2017). Further, only six of 22 QTLs had effects opposite to those expected and, for some (e.g., the flowering date effects of LH1) the locus-level effects were instead consistent with expectations of the fast-slow life history axis. This suggests that our results are broadly representative of the genetic basis *of M. cardinalis* adaptive divergence across climatic gradients.

## CONCLUSIONS

Understanding the evolutionary past and future potential of plant adaptation requires knowledge of the genetic mechanisms underlying differentiation in individual traits and correlated trait syndromes. Our genetic characterization of life-history, physiological, and mating system divergence between scarlet monkeyflower (*Mimulus cardinalis*) populations adapted to distinct seasonal habitats provide an important step toward understanding the correlated trait syndromes in this ecological model species.

First, we find that intra-specific differentiation between our parental populations (partly) parallels coordinated shifts in life-history, mating system and physiology seen at larger taxonomic scales. Second, we demonstrated high heritability for many individual traits, as well as genetic correlations potentially contributing to functional associations, but found low heritability for several key leaf economics spectrum traits (e.g., LMA). Third, by constructing a high-density gene-based intraspecific linkage map, we identified QTLs for several traits of interest, most notably a Mendelian locus controlling vegetative anthocyanin and separate multi-trait QTLs underlying early growth and flowering, late growth and asexual reproduction, and leaf thickness and gas exchange traits, respectively. The latter argue for more detailed studies of leaf anatomy variation (including direct measures of laminar thickness, cell structure and venation) across *Mimulus* populations, and broadly across herbaceous plants. Overall, these findings suggest that traits associated with a drought avoidance syndrome in the annualized southern population have a modular genetic basis, allowing for independent evolution of the different components, but that functional coordination of both life history and leaf physiological traits may allow rapid adaptation to novel climatic conditions across the species range.

## AUTHOR CONTRIBUTIONS

This study was designed by LF, AMS, and CDM, with input from ALA. AMS and CDM generated phenotypic data. LF, AMS, and TCN generated genotypic data, with assistance from DDV and KA. TCN and LF analyzed data and LF wrote paper with contributions from all co-authors.

## ACKNOWLEDGMENTS

The authors thank Findley R. Finseth and Tamara Max for assistance with data collection and members of the Angert Lab for helpful comments on previous versions of the manuscript. Funding was provided by *NSF DEB-1407333 to AS and LF, and NSF DEB-1457763 and OIA-1736249 to LF.*

## DATA AVAILABILITY

The phenotypic data will be archived on Dryad and sequencing data on the NCBI Sequence Read Archive prior to publication.

